# Playbacks of Food-associated Calls Attract Chimpanzees Towards Known Food Patches in a Captive Setting

**DOI:** 10.1101/2020.11.13.381996

**Authors:** Lisa R. O’Bryan, Susan P. Lambeth, Steven J. Schapiro, Michael L. Wilson

## Abstract

Food-associated calls have attracted much research attention due to their potential to refer to discovered food in a word-like manner. Nevertheless, their effect on receiver behavior remains unclear for many species. While some studies suggest that food-associated calls attract other foragers, other studies indicate that they repel others. We conducted playback studies to differentiate between these two hypotheses for the function of the chimpanzee (*Pan troglodytes*) food-associated ‘rough grunt’. We tested how acoustic playbacks of rough grunts (or control calls) from one of two known, identical food patches affected receivers’ foraging decisions in a captive setting. We found that participants were more likely than chance to first investigate the patch from which rough-grunts, but not control calls, were broadcast. However, neither condition increased the likelihood that participants fed first from a given patch. Our results support the hypothesis that rough-grunts attract receivers. However, since receivers were already aware of the presence of food, our results question whether rough-grunts attract by conveying information about discovered food, or rather, the signaler’s motivational state.

## Introduction

Many species of birds and mammals produce vocalizations while foraging. These ‘food-associated calls’ represent an evolutionary puzzle since drawing attention to discovered food may attract competitors to the feeding site, reducing the signaler’s food intake, and, thereby, negatively affecting the signaler’s reproductive success. Compounding this puzzle, studies have reported that food-associated calls may be capable of advertising the attractiveness of the discovered food by conveying information about its high quality or abundance. These information-sharing properties have led some researchers to label food-associated calls as ‘functionally referential signals’ (Evans & Evans, 1999; Kitzmann & Caine, 2009; Slocombe & Zuberbühler, 2005) and look toward this calling behavior for insight into the evolutionary origins of human language (Fedurek & Slocombe, 2011; Zuberbühler, 2003). Despite substantial research interest in food-associated calling behavior, how signalers benefit from producing these signals remains unclear for many species (Clay, Smith, & Blumstein, 2012). Since signals are only expected to evolve when signalers gain a net benefit from the change in behavior elicited from receivers (Krebs & Dawkins, 1984), determining the function of this calling behavior from the signaler’s perspective is key to understanding the evolutionary origins of this seemingly altruistic communication behavior.

Due to the potential for food-associated calls to alert others to the presence, and even properties, of discovered food, most research exploring the function of food-associated calling behavior has focused on its ability to attract others to the food and the benefits receivers may gain by increasing foraging group size. These studies indicate that food-associated calls can promote food searching behavior in receivers, increase the likelihood that others approach the discovered food (Chapman & Lefebvre, 1990; Elgar, 1986a; Heinrich, 1988), reduce the amount of time it takes for others to arrive at the food patch (Chapman & Lefebvre, 1990; Elgar, 1986a), and increase the total number of individuals that arrive (Brown, Bomberger Brown, & Shaffer, 1991; Laidre, 2006). By increasing foraging group size, this signaling behavior may reduce the signaler’s risk of predation or enable the signaler to reduce levels of vigilance while foraging (Chapman & Lefebvre, 1990; Elgar, 1986b), increase mating (Di Bitetti, 2005; Evans & Marler, 1994) or other social opportunities (Fedurek & Slocombe, 2013), or increase the signaler’s ability to defend the discovered food (Heinrich, 1988). For example, immature vagrant ravens (*Corvus corax*) produce food-associated calls upon discovery of a carcass in an adult male’s territory. These vocalizations have been found to attract other vagrants to the site, increasing their ability to defend the carcass from the territory-holder (Heinrich, 1988). As another example, roosters (*Gallus gallus domesticus*) produce food-associated calls when hens are nearby, prompting hens to approach and consume the food while increasing the likelihood that she mates with the signaler (Pizzari, 2003). In addition, house sparrows (*Passer domesticus L.*) are more likely to produce food-associated calls upon discovery of a food patch when they are alone or in a small group and when the patch is far from cover. This calling behavior increases the size of the foraging group, presumably diluting the signaler’s risk of predation (Elgar, 1986b). Thus, even though food-associated calling behavior has the potential to increase feeding competition for the signaler, the net benefit gained by increasing foraging group size or neighbor proximity may promote the evolution of this calling behavior in many species.

Despite the strong research focus on the attractive nature of food-associated vocalizations, several studies provide evidence that some vocalizations may have the opposite function from the signaler’s perspective. For example, Radford and Ridley (2008) found that the food-associated calls of pied babblers (*Turdoides bicolor*) do not attract others toward the signaler, but rather reduce the likelihood that others approach the signaler while foraging. As another example, food-associated calling behavior in white-faced capuchins (*Cebus capucinus*) has been found to correlate with an increase in spacing between individuals already foraging within the patch (Boinski & Campbell, 1996) and a reduction in the likelihood that the signaler is closely approached, or receives aggression, by other foragers (Gros-Louis, 2004a). In addition, several studies have found evidence that the food-associated calling behavior of some species positively correlates with the signaler’s hunger level (Hauser & Marler, 1993; Heinrich & Marzluff, 1991), its motivation to feed in a particular location (Fedurek & Slocombe, 2013) and/or its high dominance status (Clark & Wrangham, 1994; Heinrich & Marzluff, 1991). These studies suggest that vocalizations produced in foraging contexts do not necessarily attract others to the food. Rather, some may have the opposite effect-repelling others by advertising the signaler’s ownership of, motivation for, or willingness to defend, the food.

Recently, several researchers have focused their attention on the food-associated ‘rough grunt’ of chimpanzees (*Pan troglodytes*). While the term ‘rough grunt’ implies a single discrete vocal category, acoustic analyses indicate that this term includes a set of graded signals that range from noisy, low-pitched grunts to tonal, high-pitched barks (Slocombe & Zuberbühler, 2006) with calling bouts that can vary from one to many vocalizations (Brosnan & de Waal, 2000; Fedurek & Slocombe, 2013). In contrast to the chimpanzee pant hoot vocalization, which may be produced not only upon arrival at a food patch, but also in a variety of other contexts (Clark & Wrangham, 1994), rough grunts are only produced in foraging contexts (Goodall, 1986). Because of this contextual specificity, this vocalization has been labeled a functionally referential signal (Slocombe & Zuberbühler, 2005), and several studies have explored its ability to provide others with information about discovered food. These studies suggest that variation in rough grunt acoustic properties may convey information about the quality (Fedurek & Slocombe, 2013; Slocombe & Zuberbühler, 2006), divisibility (Brosnan & de Waal, 2000; Hauser, Teixidor, Fields, & Flaherty, 1993), or even identity of some foods (Slocombe & Zuberbühler, 2005). Accordingly, some researchers have proposed that rough grunts play an important role in chimpanzee societies by facilitating the sharing of food with group members (Brosnan & de Waal, 2000; Fedurek & Slocombe, 2013).

Most prior research on rough grunt calling behavior has focused on the context of call production and the information conveyed by these calls. In contrast, only a few studies have examined the effect these vocalizations have on receiver behavior or how this change in behavior may affect signalers. The only prior experimental study on this topic suggests that differences in rough grunt acoustics can inform listeners about the availability of specific foods, enabling receivers to adjust the location and intensity of their foraging effort according to their preference for that food. (Slocombe & Zuberbühler, 2005). However, as this study was only conducted with a single individual, it is unclear whether these results are generalizable to other individuals and/or contexts. Notably, even though several studies conducted in captivity have reported a correlation between properties of discovered food and the production and acoustic properties of rough grunts (Brosnan & de Waal, 2000; Hauser et al., 1993; Slocombe & Zuberbühler, 2005, 2006), studies conducted in the field have largely failed to replicate these findings (Kalan, Mundry, & Boesch, 2015; Slocombe & Zuberbühler, 2006) (though see (Fedurek & Slocombe, 2013)). A few observational studies have examined the effect that rough-grunts have on receiver behavior. One study found that rough-grunts did not correlate with the subsequent arrival of other foragers in a food patch, except for when considering feeding bouts involving fruit (Kalan et al., 2015). Another study similarly found that the production of rough-grunts upon arrival only correlated with the arrival of more extra-party individuals when considering one tree species. An additional study found that rough-grunt production was correlated with an increased likelihood that close associates remained nearby while the signaler fed (Fedurek & Slocombe, 2013). However, this study did not examine whether these individuals shared the same food patch as the signaler. Thus, evidence that rough-grunts attract receivers to the signaler’s food patch remains tentative.

Social context has been found to play a central role in rough grunt production. Chimpanzees produce rough grunts more often when foraging with others than when alone (Brosnan & de Waal, 2000; Slocombe et al., 2010), and Schel et al. (2013) found that chimpanzees that were previously feeding silently began producing rough grunts only once the researchers used acoustic playbacks to simulate the arrival of another chimpanzee. Chimpanzees are especially more likely to produce rough-grunts when their current party includes higher ranking individuals (Schel et al., 2013) or close associates (Fedurek & Slocombe, 2013; Schel et al., 2013; Slocombe et al., 2010) and they are more likely to produce rough-grunts after an aggressive event had occurred in a food patch (Ischer, Zuberbühler, & Fedurek, 2020). In addition, chimpanzees have been found to produce rough-grunts when they, themselves, are motivated to forage in a given patch (Fedurek & Slocombe, 2013). Thus, there is high potential that rough-grunts provide socially relevant motivational information to nearby foragers, rather than, or in addition to, information about food.

In an effort to clarify the function of rough grunt vocalizations, we sought to test the assumption that rough grunts are attractive vocalizations while also testing the opposite hypothesis that rough grunts are repulsive to receivers. We conducted a series of playback studies with captive chimpanzees to test how playbacks of rough grunt vocalizations affected receivers’ decisions to forage on one of two known, identical food sources. Participants had the chance to approach either a silent food patch or one from which the vocalizations of a familiar group member had been broadcast. By providing participants with the silent food patch in addition to the patch from which the stimulus was broadcast, we aimed to reduce the likelihood that participants would approach the stimulus simply because it was their only option for acquiring food. We believed this was important because some studies have found that individuals may be attracted to food-associated calling behavior that is locally repulsive simply because its contextual specificity provides information of a potential feeding opportunity (Gros-Louis, 2004b; Heinrich & Marzluff, 1991). We predicted that, if rough grunts attract others, participants would be drawn to the food patch from which rough grunts were broadcast. However, if rough grunts repel others, we predicted that participants would first approach the food patch from which these vocalizations had not been broadcast. Lastly, if rough grunts neither attract nor repel, we predicted that participants would be equally likely to approach either food patch since they were already aware of the presence and identical quality of the two food sources.

## Material and methods

### Study site and participants

L.O’B. (‘the observer’) collected data from June-August 2010 from chimpanzees housed at the National Center for Chimpanzee Care at the Michale E. Keeling Center for Comparative Medicine and Research. This site houses a large population of chimpanzees that reside in approximately 20 multi-male, multi-female social groups. As part of their normal living conditions, all animals involved in our study had *ad libitum* access to an indoor and outdoor enclosure, as well as to monkey chow and water. In addition, individuals were fed four fresh produce meals per day and participated in food- and/or drink-related enrichment activities multiple times each week. 12 adult chimpanzees (7 female and 5 male, Table 1) from two different social groups **voluntarily** participated in our study. These participants were not deprived of food or water at any time during the study and had access to chow and water during all trials. While participants did occasionally drink water during trials, none consumed chow.

**Table 1.** Basic demographic information for study participants

| Participant ID | Group ID | Age | Sex | Birth and Rearing Environment |
| --- | --- | --- | --- | --- |
| BD | 1 | 32 | Female | Captive Born Nursery Reared |
| JD | 1 | 20 | Male | Captive Born Nursery Reared |
| KB | 1 | 38* | Male | Wild Born |
| PT | 1 | 41* | Female | Wild Born |
| TK | 1 | 30 | Female | Captive Born Mother Reared |
| QY | 1 | 39* | Female | Wild Born |
| BK | 2 | 25 | Female | Captive Born Nursery Reared |
| GI | 2 | 27 | Male | Captive Born Mother Reared |
| KK | 2 | 26 | Male | Captive Born Mother Reared |
| KP | 2 | 19 | Female | Captive Born Mother Reared |
| NO | 2 | 24 | Male | Captive Born Mother Reared |
| TA | 2 | 21 | Female | Captive Born Mother Reared |

Data on each participant's age and rearing environment were obtained from records kept by staff members at KCCMR. Asterisks following the ages of the wild-born participants indicate that their ages are estimated since their exact years of birth are not known. Age is calculated with reference to the start of our study in June 2010.

### Set-up

The study environment consisted of each social group’s three adjacent indoor rooms positioned in a row (Figure 1). Each room had a sliding door separating it and its adjacent room(s), as well as the outdoor area. All doors could be manipulated by the observer from outside of the chimpanzee area (in the ‘human area’). The walls separating the chimpanzee and human areas were largely made of wire mesh, enabling the observation and documentation of the participants’ behavior from the human area. One video camera on a tripod was positioned in the human area outside of each of the three rooms, focusing on the center of each room. This enabled the participant’s activity to be documented as it traveled in and between each room. The walls between each of the three rooms were opaque so that participants could not see the insides of each room without looking through an open door or entering the room.

**Figure 1.**
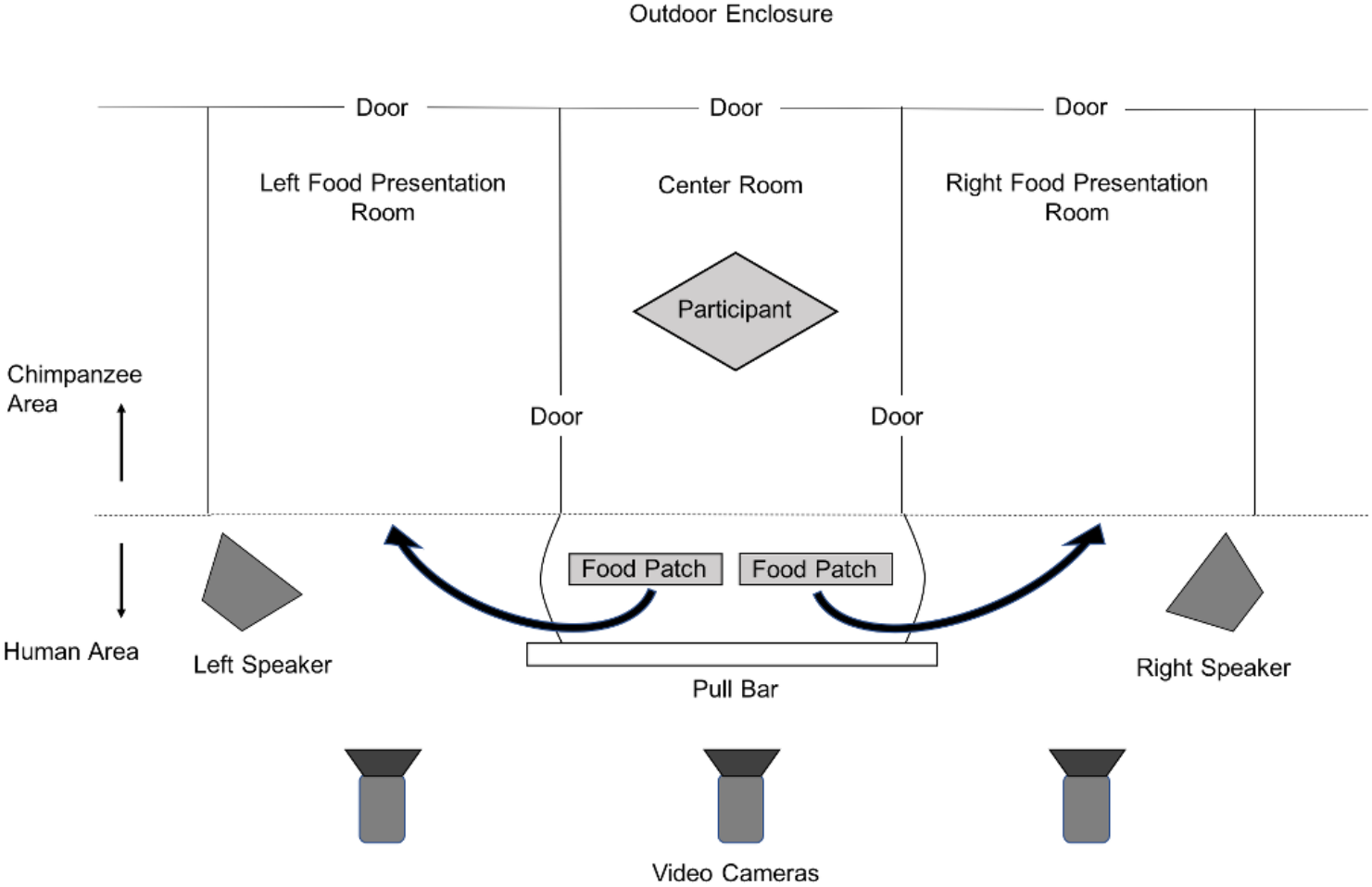
The set-up as it looked just after the participant voluntarily entered the study area from the outdoor enclosure. Note that the figure is not drawn to scale. The arrows indicate where the observer placed the food patches prior to the initiation of playback. The pull bar was used to simultaneously open both doors between the center room and food presentation rooms, allowing the participant to enter.

Each food patch consisted of 60 grapes resting in a trough composed of a polyvinyl chloride (PVC) pipe that had been cut in half lengthwise. We chose grapes as the food item in this study because food choice tests conducted by Hopper et al. (2013) indicated that they are a preferred food in this population of chimpanzees. Similar to other enrichment devices used to engage chimpanzees at this study location, we attached the food patches to the outside (i.e. human side) of the wire mesh between the human and chimpanzee areas so that they could be readily attached and detached by the observer. Participants consumed the grapes by sticking their fingers through the mesh and pulling the grapes through the openings.

We broadcast vocalizations to participants through one of two speakers (Mackie SRM 350v2) placed in the human area just outside of each side room and angled toward the center room (Figure 1). The center of each speaker was positioned approximately at the height of a chimpanzee’s head while feeding at the food patch. The speakers could not be seen by the participants from the center room, but could be viewed once the participant entered the food presentation rooms. Since we never broadcast vocalizations when the participant was in view of the speakers, and since the speakers were within view even during trials when no vocalizations were broadcast, we did not expect participants to associate these speakers with the vocalization playbacks. A laptop containing the stimulus sound files was placed in the far corner of the human area.

### Procedure

All participants experienced two training trials and three study trials. The training trials were carried out in the same manner as the study trials except that no stimuli of any kind were presented to the participants. We conducted the training trials to familiarize the participants with 1) the study procedure and 2) the fact that they were able to approach and feed from both food patches during each trial.

Participants engaged in no more than one trial per day. At the start of each trial, all doors in the study area were in the closed position. Participants voluntarily entered the study area when the door between the outdoor enclosure and the center room was opened. Once the participant was inside, the observer closed the door behind the chimpanzee. If a participant was not entirely comfortable while in the center room, or at any other time during the study, the observer was capable of opening the door to the outdoor enclosure so that the chimpanzee could choose to go back outside. At the time of the participant’s entry, the two food troughs were lying next to one another on the ground in the human area just outside of the center room, so that the participants could observe the presence and equal quality of both food sources (Figure 1). Once the door was closed, the observer attached one food trough to the wire mesh of one of the side rooms. The observer then attached the other trough to the other room in the same manner. We randomly selected the side to which a trough was first attached before each trial. During the study trials, the observer initiated the playback once the food was placed.

There were three treatment levels: ‘Silence’, ‘Rough Grunt’ and ‘Control Call’. Each participant experienced all three treatment levels in a randomized order. For a given participant, the identity of the individual producing the rough grunt and control call was kept consistent. The Silence condition was used to determine whether participants had a significant side bias. The Rough Grunt condition was used to determine whether, and how, rough grunts influenced the participants’ foraging decisions. The Control Call condition controlled for the presence and vocal activity of a specific individual, enabling us to determine whether the participants were responding to the rough grunts or simply the presence of another individual.

During both the Rough Grunt and Control Call conditions, a rough grunt or control call, respectively, was broadcast from one of the two speakers while a silent stimulus was broadcast from the other. We randomly selected the side from which the rough grunt or control call was broadcast before each trial. The observer initiated the playback by discretely pressing play on a remote control while standing in the human area in line with the middle of the center room. Once the playback was completed, the observer simultaneously opened the sliding doors separating the center room from the two food presentation rooms using a pull bar that was connected on each end to each of the two doors (Figure 1). Prior to this point, participants were not capable of viewing the contents of either room. Once the doors were opened, participants were capable of freely investigating both rooms. Once all of the food was consumed from both rooms, the observer opened a door to the outdoor area so that the participant could exit the study area. The Silence condition was carried out in the same manner as the Rough Grunt and Control Call conditions, except that silent stimuli were broadcast from both speakers.

### Playback stimuli

L.O’B. recorded all playback stimuli with a Sennheiser ME66 shotgun microphone with K6 power module and a Marantz PMD670 recorder. Most vocalizations were recorded *ad libitum* during regular mealtimes and/or social interactions within the group. However, some rough grunts were elicited by placing food (grapes or other produce) inside the chimpanzees’ enclosures. While the possibility exists that the acoustic properties of rough grunts convey information about food quality or even food type (Brosnan & de Waal, 2000; Fedurek & Slocombe, 2013; Hauser et al., 1993; Kalan et al., 2015; Slocombe & Zuberbühler, 2005, 2006), here we focus on the question of whether the calls are generally attractive or repulsive. Therefore, we do not expect that the specific stimulus that elicited the calls should affect behavior at this level. Stimuli were recorded from four different individuals, two from each social group. One rough grunt and one control call stimulus was created from the recordings obtained from each of the four individuals (Table 2). Due to the limited time frame of the study, we were not able to record enough vocalizations from each individual to permit playback of a single call type for all Control Call trials. Instead, the type of vocalization presented to participants during the Control Call condition varied across participants. Nevertheless, these vocalizations controlled for the fact that the participant heard the vocalization of another chimpanzee in the adjacent room. Two of the control calls used in the study were ‘pant-grunts’: calls produced by lower ranking chimpanzees to higher ranking chimpanzees (Bygott, 1979; Goodall, 1986). One of the control calls was a ‘pant hoot’: a long-distance call produced in a variety of contexts, including a foraging context (Reynolds & Reynolds, 1965; Wrangham, 1977). The other control call was a ‘raspberry’: a vocalization frequently produced in captive environments to catch the attention of humans (Hopkins, Tagliatela, & Leavens, 2007). We presented three participants from each group with one of the four unique Rough Grunt/Control Call stimulus pairs (Table 2). We left stimuli unedited except for reducing the stimulus length to a duration of 6 seconds. We cropped the stimulus files using Praat Version 5.1.34 (www.praat.org) and broadcast the signals using Windows Media Player (version 11, © 2006 Microsoft Corporation).

**Table 2.** Playback stimulus information

| Group ID | Signaler | Receivers | Control Call Type |
| --- | --- | --- | --- |
| 1 | JD | BD, PT, KB | Raspberry |
| 1 | BD | JD, TK, QY | Pant grunt |
| 2 | KP | BK, GI, NO | Pant grunt |
| 2 | RE | KK, KP, TA | Pant hoot |

A rough grunt was also recorded by all signalers for use in the Rough Grunt condition. The identity of the signaler was held consistent for the Rough Grunt and Control Call conditions

### Analysis

We focused our analysis on which room the participants investigated first during the study trials. A participant was considered to investigate a room if s/he walked towards the door of a room and peered inside and/or immediately entered the room. We also analyzed which room the participants fed in first, since participants did not always proceed to feed from the room they first investigated (Supplemental Figure 1). A participant was considered to feed in a given room if s/he consumed any food from that food patch. For the silent condition, we used chi-squared tests to determine whether the number of participants that investigated and fed first in the left (vs. right) room differed significantly from chance. For the rough grunt and control call conditions, we used chi-squared tests to determine whether the number of participants that investigated and fed first in the Stimulus (vs. Non-stimulus) Room differed significantly from chance. For a given trial in the Rough Grunt and Control Call conditions, the ‘Stimulus Room’ was considered to be the food presentation room from which a chimpanzee vocalization was broadcast, while the ‘Non-stimulus Room’ was the food presentation room from which a silent stimulus was broadcast.

## Results

All participants voluntarily and successfully completed all training and study trials. During a Control Call trial, participant ‘KK’, an adult male (Table 1), produced a pant hoot in response to the playback of a pant hoot by ‘RE’, another adult male in his group. This ad lib observation suggests that the participants interpreted the acoustic stimuli as they would calls produced spontaneously by their group members. In the Silent condition, participants were equally likely to investigate and feed first in either room (Investigate: Left:Right=5:7, χ_1_2= 0.33, P = 0.56; Feed: L:R=6:6, χ_1_2= 0, P = 1.00; Figure 2). In the Rough Grunt condition, participants were more likely to first investigate the room from which a rough grunt had been broadcast compared to the silent room (Stimulus(S):Non-stimulus(NS)=10:2, χ_1_2= 5.33, P = 0.02; Figure 2a). However, participants were not more likely to first investigate the room from which a control call had been broadcast (S:NS=7:5, χ_1_2= 0.33, P = 0.56; Figure 2a). In both the Rough Grunt and Control Call conditions, participants were equally likely to feed first from either room (Rough Grunt: S:NS=6:6, χ_1_2= 0, P = 1.00; Control Call: S:NS=8:4, χ_1_2= 1.33, P = 0.25; Figure 2b). In summary, we found that participants were more likely to first approach the food patch from which rough grunts were broadcast, rather than the silent food patch, but that this preference for the Stimulus Room was not observed when control calls were broadcast. Neither call type increased the likelihood that a participant would feed first from a given food patch. All data analyzed during this study are included in Table 3.

**Figure 2.**
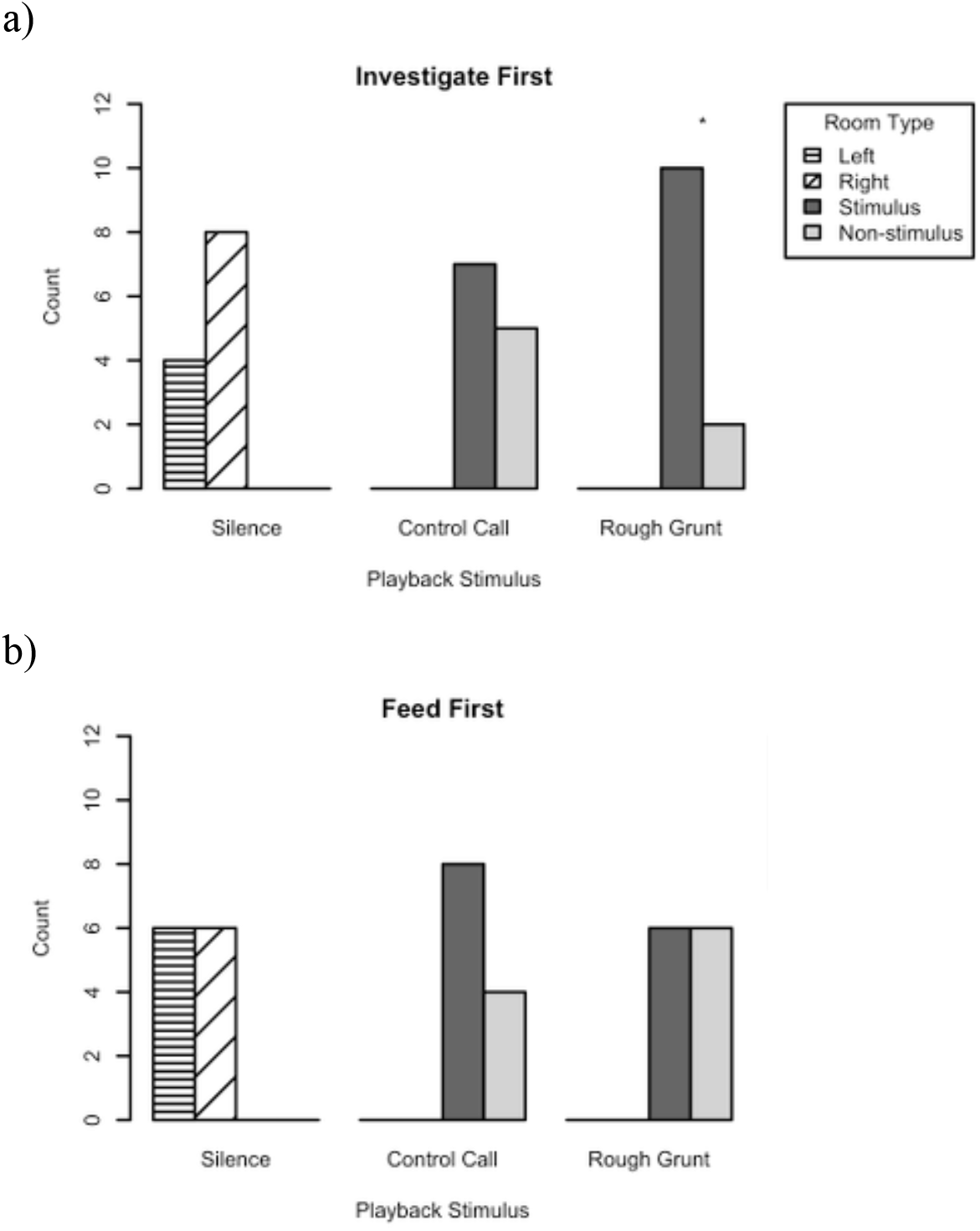
Bar plot displaying the number of participants that chose to first a) investigate and b) feed in each room in each condition. The bars for the Rough Grunt and Control Call conditions represent the number of participants that visited the Stimulus vs. Non-stimulus rooms, respectively. The bars for the Silence trial type represent the number of individuals that visited the left vs. right room, respectively. The asterisk indicates that the number of participants that first investigated the Stimulus vs. Non-stimulus room during the Rough Grunt condition differed significantly from chance (p < 0.05).

## Discussion

In this study, we sought to test whether rough grunts attract, repel or have no effect on the proximity of receivers in foraging contexts. We found that, more often than by chance, participants first approached the food patch from which rough grunts were broadcast, rather than the silent food patch. This approach behavior was not simply due to the perception that a familiar group member was present in the Stimulus Room, since the vocalizations presented in Control Call trials did not attract the participants more than by chance. These findings suggest that rough grunt production elicits an approach response from receivers, which is in alignment with previous interpretations of rough grunt calling behavior (Goodall, 1986; Slocombe & Zuberbühler, 2005), as well as studies of food-associated calling behavior in many other species (for example (Elgar, 1986b; Pizzari, 2003)). Nevertheless, it is important to test this assumption. For instance, Wrangham (1977) initially proposed that the chimpanzee food-associated pant hoot attracted others to the feeding site, since pant hoot production was found to positively correlate with the subsequent arrival of estrous females at the food patch. However, a later study did not find a correlation between pant hoot production and the arrival of any extra-party individuals (Clark & Wrangham, 1994). Furthermore, researchers have not found acoustic differences between food-associated pant hoots and those produced in other contexts (Clark & Wrangham, 1994; Notman & Rendall, 2005). Rather, Clark and Wrangham (1994) found a correlation between pant hoot production and the presence of high-ranking individuals in the foraging party, leading them to suggest that this call advertises the signaler’s high dominance rank, rather than information about discovered food (Clark & Wrangham, 1994). While there remains some uncertainty regarding the function of the pant hoot vocalization, these studies demonstrate the importance of testing assumptions about the function of food-associated, and other, vocalizations.

The results of our study did not provide any support for the hypothesis that rough grunts repel receivers. If rough grunts repel others, we predicted that receivers would first approach the silent food patch during the Rough Grunt condition. Rather, we found the opposite response. While studies have found that some receivers will approach repellant food-associated calls since they may suggest the presence of a novel food source (Gros-Louis, 2004b; Heinrich & Marzluff, 1991), it is unlikely that participants approached the Stimulus Room because they believed it was their only chance for acquiring food. This is because we designed our study so that participants had the choice of feeding in either the Stimulus or Non-stimulus Room.

Researchers have commonly assumed that rough grunts elicit an approach response in receivers due to the information they provide about the presence and/or properties of the food source (Clay et al., 2012; Goodall, 1986; Kalan et al., 2015; Slocombe & Zuberbühler, 2005, 2006). However, it is unlikely that this was the reason for the attractive response we observed in our participants. This is because we familiarized the participants with the presence and equal quality of both food patches during the training trials, and, at the start of each trial, participants had the opportunity to view both food patches before the observer attached them to the outside of each food presentation room (Figure 1). Thus, it is unlikely that the participants approached the Stimulus Room because they believed that this food patch was of higher quality than the other, especially since they were not more likely to feed first from this food patch. However, it is possible that participants associated the rough grunt playbacks with the presence of an additional food source since they were not able to view the inside of either food presentation room before or during the playback. In order to rule out this possibility, a follow-up study similar to that conducted by Slocombe and Zuberbuhler (2005) would need to be conducted in order to assess whether chimpanzees have specific expectations about the presence and properties of a food source upon hearing specific rough grunt vocalizations.

An alternative reason for why rough grunts attract listeners may be that they advertise the signaler’s motivation to forage for a long period of time in a particular location-information that could help promote behavioral coordination between group members. In a study testing this hypothesis, Fedurek and Slocombe (2013) found that rough grunt production by wild chimpanzees was correlated with longer feeding bouts by the signaler. Furthermore, they found that an adult male’s important social partners were more likely to remain in the vicinity of the food patch until he finished feeding when he produced rough grunts compared to when he did not. Also, chimpanzees that remained silent while foraging were found to feed longer when others in their party produced rough grunts (Fedurek & Slocombe, 2013; Slocombe & Zuberbühler, 2006). Thus, rough grunt production could help signalers retain important group members in the vicinity of the food patch, since chimpanzees (Georgiev et al., 2014), like other species (Alberts, Altmann, & Wilson, 1996; Kazahari, 2014), can experience a tradeoff between maximizing foraging efficiency and maintaining social cohesion.

Another possibility for why rough grunts elicit an attractive response could be that rough grunts advertise an affiliative motivational state to fellow foragers, facilitating co-feeding at food patches. Chimpanzees display high levels of food-related aggression in foraging contexts (Goodall, 1986; Wittig & Boesch, 2003). Therefore, the production of food-associated calls could signal when it is safe for others to approach and co-feed, such as when food is abundant (Brosnan & de Waal, 2000; Fedurek, Donnellan, & Slocombe, 2014). Encouraging others to cofeed at an abundant food patch could be beneficial for both signaler and receiver, since it could reduce the risk of predation (Elgar, 1986b), deter others from seizing the food patch (Heinrich, 1988) and/or promote opportunities that may mutually enhance fitness, such as mating or grooming. Similarly, by advertising a reduced likelihood of challenging others, signalers could reduce the chances that they, themselves, will receive aggression, as was observed in white-faced capuchins (Gros-Louis, 2004a). This hypothesis, that rough grunts advertise an affiliative motivational state while foraging, is congruent with previous findings that chimpanzees are more likely to produce rough grunts in the presence of both close social partners and high-ranking individuals and after the occurrence of aggression (Fedurek & Slocombe, 2013; Ischer et al., 2020; Schel et al., 2013).

In summary, we found evidence that rough grunts elicit an approach response from receivers, supporting the hypothesis that these vocalizations are attractive rather than repulsive. This finding is important because it helps to clarify the function of rough grunt vocalizations. Since the vast majority of research on rough grunt vocalizations has focused on their potential ability to inform others about discovered food (for a recent review see: (Clay et al., 2012)), more studies are needed to clarify the effect these vocalizations have on the behavior of receivers, as well as how this change in behavior benefits signalers. We suggest that the findings of this study should be tested with wild chimpanzees by studying the association between rough grunt production and changes in neighbor proximity and/or co-feeding behavior. Furthermore, in order to test the hypothesis that rough grunts advertise an affiliative motivational state, studies could investigate the correlation between rough grunt production and the likelihood of aggression by signalers or the likelihood that signalers receive aggression by others. By more thoroughly investigating aspects of food-associated calling behavior, other than its potential to function referentially, we can gain a more rounded view of this widespread vocal behavior and gain a better understanding of its function (Owren & Rendall, 2001).

## Acknowledgements

We would like to thank all of the staff at the National Center for Chimpanzee Care for their wonderful care of the chimpanzees housed at the facility. LO’B would also like to give special thanks to staff members Rachel Haller, Emily Mocarski and Erica Thiele for their assistance with logistical aspects of the study and to Dr. Lydia Hopper for her helpful guidance and feedback throughout the study.

## Ethical Approval

All applicable international, national, and/or institutional guidelines for the care and use of animals were followed. All procedures performed in studies involving animals were in accordance with the ethical standards of the institution at which the studies were conducted and were approved by the Institutional Animal Care and Use Committee (IACUC) of the University of Minnesota (Protocol number #1002A78194). This article does not contain any studies with human participants performed by any of the authors.

