## Supplemental Figure 1 for "Playbacks of Food-associated Calls Attract Chimpanzees Towards Known Food Patches in a Captive Setting"

**Supplemental Figure 1**: Response of participants in each playback condition

| ID | Stimulus Type | Trial Order | Stimulus Side | Investigated First | Fed First |
| --- | --- | --- | --- | --- | --- |
| BD | Control | 1 | Left | Right | Right |
|  | RG | 3 | Right | Right | Right |
|  | Silence | 2 | NA | Right | Right |
| PT | Control | 2 | Left | Left | Left |
|  | RG | 1 | Right | Right | Left |
|  | Silence | 3 | NA | Left | Left |
| KB | Control | 2 | Right | Left | Left |
|  | RG | 3 | Right | Right | Right |
|  | Silence | 1 | NA | Left | Left |
| BK | Control | 3 | Right | Right | Right |
|  | RG | 2 | Right | Right | Right |
|  | Silence | 1 | NA | Right | Right |
| GI | Control | 1 | Right | Right | Right |
|  | RG | 3 | Right | Right | Right |
|  | Silence | 2 | NA | Right | Right |
| NO | Control | 2 | Left | Right | Right |
|  | RG | 3 | Right | Right | Right |
|  | Silence | 1 | NA | Right | Left |
| JD | Control | 1 | Right | Right | Right |
|  | RG | 3 | Right | Right | Left |
|  | Silence | 2 | NA | Right | Right |
| TK | Control | 3 | Left | Left | Left |
|  | RG | 1 | Left | Left | Left |
|  | Silence | 2 | NA | Left | Left |
| QY | Control | 3 | Left | Left | Right |
|  | RG | 2 | Right | Left | Left |
|  | Silence | 1 | NA | Left | Right |
| KK | Control | 2 | Left | Left | Left |
|  | RG | 1 | Left | Right | Right |
|  | Silence | 3 | NA | Right | Left |
| KP | Control | 3 | Left | Right | Left |
|  | RG | 1 | Left | Left | Right |
|  | Silence | 2 | NA | Right | Left |
| TA | Control | 1 | Left | Right | Left |
|  | RG | 2 | Left | Left | Right |
|  | Silence | 3 | NA | Right | Right |

^The ‘Stimulus side’ column indicates whether a given stimulus was broadcast from the left or right food presentation room in the Rough Grunt and Control Call conditions.^
